# A Way Through the Trees: Molecular Phylogenies Consistently Recover Two Clades of *Aedes* Mosquitoes

**DOI:** 10.1101/2025.07.16.665186

**Authors:** Huiqing Yeo, Carmen Chan, Gen Morinaga, Brian M. Wiegmann, John Soghigian

**Author notes:** These authors contributed equally to this manuscript (Huiqing Yeo, Carmen Chan).

## Abstract

Phylogenetic analyses of molecular data have been critical in resolving deep and shallow relationships across the tree of life. Such analyses have provided clarity where traditional morphological analyses may not have, and have been instrumental in a range of evolutionary, systematic, and taxonomic studies for providing clarity to evolutionary relationships. One such case where evolutionary relationships are poorly understood is in the mosquito genus *Aedes*. Although this medically important genus of insects has received significant study, the majority of research has focused on those relatively few species that are invasive and transmit pathogens which cause disease. As a result, evolutionary relationships between Aedes mosquitoes are poorly understood, and the classification of the genus has been contentious, with numerous taxonomic changes over the last two decades being undertaken and reversed based entirely on analyses of somewhat limited morphological data. As a result, we conducted a literature review of molecular phylogenies and evaluated classificatory hypotheses under a framework based on observations of frequently occurring relationships among mosquito genera and subgenera. We found that molecular phylogenies consistently reflect two distinct clades of *Aedes* mosquitoes, each with other Aedini genera within them. We discuss these results in the context of ongoing debates over *Aedes* nomenclature. Our review demonstrates the ability for molecular phylogenies to aid in resolving long-standing debates on nomenclature.

**Highlights:**

- The genus Aedes, in the tribe Aedini, contains many of the world’s most important disease vector mosquitoes, such as the yellow fever mosquito Aedes aegypti.
- The phylogeny of the genus Aedes remains contentious, leading to repeated and often confusing changes in taxonomy and nomenclature.
- No comprehensive molecular phylogenies of the genus Aedes have been published the, but Aedes species have been included in many phylogenetic analyses.
- Molecular phylogenies demonstrate consistent results: There are two major clades of *Aedes* mosquitoes, neither of which are monophyletic.

**Graphical Abstract:** 

## 1 Introduction

Phylogenetic analyses of molecular data have been critical in resolving the deep and shallow patterns of evolutionary relationships across the tree of life. In insects, this has been particularly profound, resulting in major new classificatory insights such as the ‘death’ of an order (Inward et al., 2007) and a revised and foundational understanding of the early branching events in the insect tree of life (Misof et al., 2014). Molecular phylogenetics have enabled the re-classification of numerous lineages, resolving long-standing disagreements found through traditional cladistic analyses based on morphology alone (Niehuis et al., 2012; Song et al., 2024; Thomas et al., 2013; Vasilikopoulos et al., 2020).

One such case of long-standing classificatory instability is in the genus *Aedes* and the tribe Aedini (Diptera: Culicidae). In traditional classifications, the genus *Aedes* included over 800 species (Wilkerson et al., 2015), and was one of ten genera in the tribe. Species within the genus *Aedes* serve as vectors for medically and veterinary important diseases (Yee et al., 2022) such as chikungunya, dengue, and yellow fever (Leta et al., 2018). The ability for some *Aedes* species to utilize artificial containers for immature stages has contributed to their increasing global prevalence and medical significance (Benedict et al., 2007; Kampen and Werner, 2014; Powell and Tabachnick, 2013). As such, extensive research has been conducted on medically important *Aedes* species such as *Aedes aegypti* and *Aedes albopictus*, and this genus contains some of the highest quality genomic resources available in insects (Chen et al., 2015; Waterhouse et al., 2008). However, the majority of research has focused on specific disease vector species, while the genus as a whole—and the Aedini, the tribe to which it belongs—has received relatively little attention, particularly in terms of molecular phylogenetics.

Over the last three decades, the genus *Aedes* has undergone numerous controversial taxonomic revisions (Reinert et al., 2009; Savage, 2005; Wilkerson et al., 2015). From 2000–2009, the genus *Aedes* was reduced from 800 species to only 12; 74 new genera were established, primarily based on the elevation of subgenera to generic status (Reinert et al., 2009, 2004). These revisions were based on cladistic analyses of morphological character data, and resulted in generic and species epithets repeatedly changing in rapid succession. For instance, *Aedes japonicus* Theobald —a major invasive vector of Japanese encephalitis virus (JEV)—was classified as *Aedes* prior to 2000, reclassified in 2000 by Reinert to *Ochlerotatus japonicus*, then subsequently reclassified again in 2006 to *Hulecoeteomyia japonica*. These changes were reverted in 2015 by Wilkerson et al. (2015) based on a re-analysis of morphological data. For the most part, these authors did not consider results from molecular phylogenies in their return to a reunified *Aedes* (and thus a 10-genus Aedini) with more than 800 species. Such rapid reclassification was met with varying degrees of acceptance (Polaszek, 2006; Rattanarithikul et al., 2010; Reisen, 2016; Savage, 2005) and public repositories still reflect the multiple (and at times conflicting) names used for *Aedes* species between 2000 and 2015; for instance, some members of the same subgenus are split between generic classifications—*Aedes togoi* (Theobald, 1907) (GenBank: *Aedes togoi*) and *Aedes savoryi* Bohart 1957 (GenBank: *Tanakaius savoryi*) both belong to the subgenus *Tanakaius* in the genus *Aedes* in the present taxonomy, but not in various public repositories (Global Biodiversity Information Facility, 2025; National Center for Biotechnology Information, 2025). This lack of standardization and stable classification can be confusing, leading to difficulties in literature search and data management and disrupting communication among workers of mosquitoes.

Although the majority of controversial taxonomic actions described above occurred in the early 2000s, the lack of monophyly in the genus *Aedes* has been recognized since at least Belkin in 1962 (Belkin, 1962), wherein he acknowledged there were two clades of *Aedes* mosquitoes: Section A, corresponding to members of the subgenera *Ochlerotatus, Mucidus, Finlaya*, the genus *Opifex*, and other subgenera; and Section B, including the subgenera *Aedes, Stegomyia, Aedimorphus, Verrallina* (now considered a separate genus), the genus *Armigeres*, and others (Figure 1A). Although his consideration included only those mosquitoes found in the South Pacific, Belkin’s conclusions were remarkable in that they captured similar groups of subgenera as those in cladistic analyses used to divide the genus *Aedes* into 74 genera four decades later (Figure 1B)—and, as we will show, they bear a striking resemblance to the groups of subgenera that are reflected in molecular phylogenies of the Aedini (Figure 1C).

**Figure 1.**
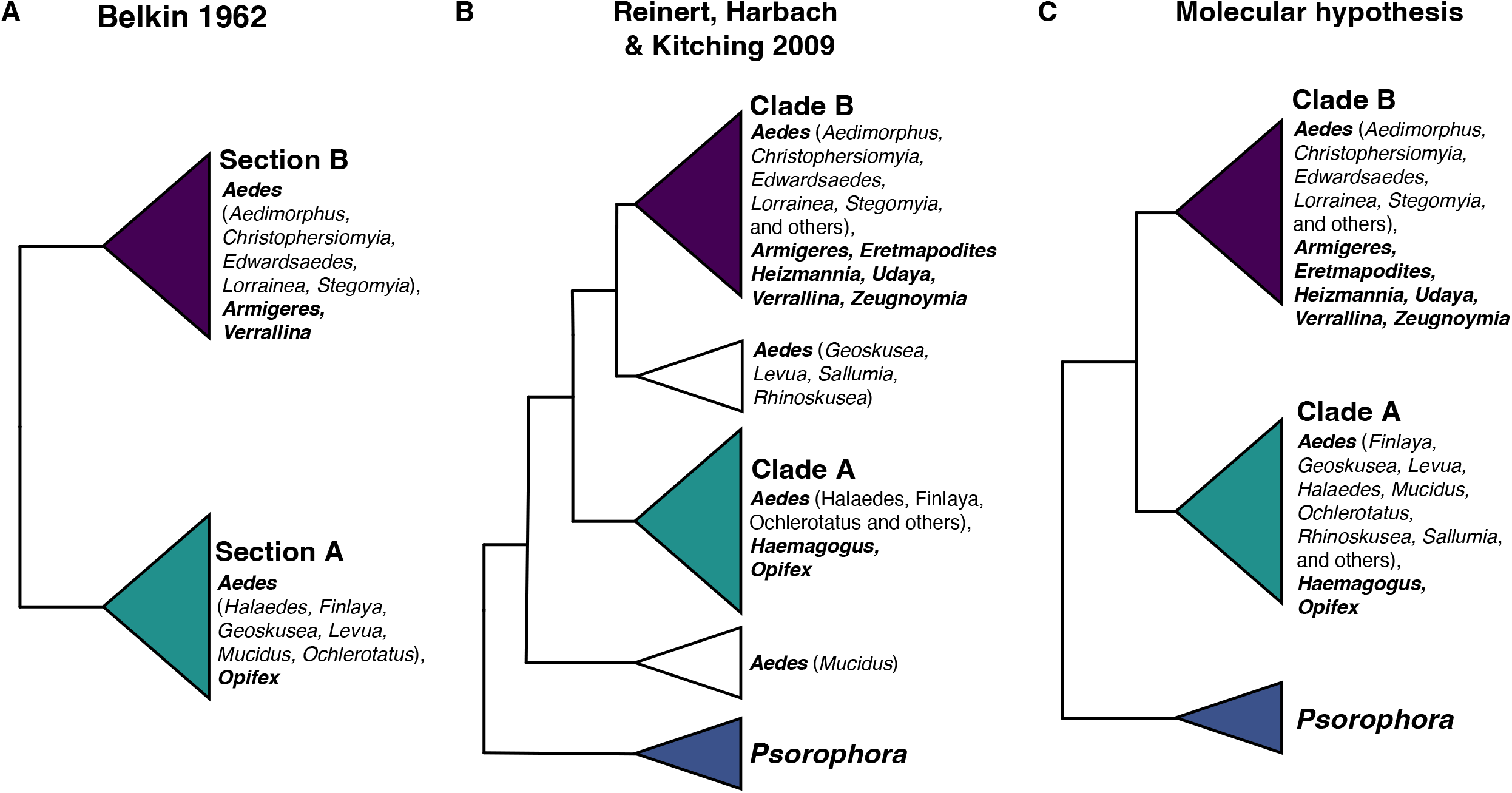
Formulation of the three-clade hypothesis for tribe Aedini. (A) In an early attempt to group aedine taxa, Belkin hypothesized that the genus *Aedes* was non-monophyletic, and that aedine taxa was split into two sections. At that time, he was working on mosquitoes of the South Pacific and did not include other genera (including *Psorophora*) and subgenera in his hypothesis. (B) Reinert, Harbach and Kitching (2009) published the largest aedine phylogeny based on morphological characters of all life stages, placing all genera and most *Aedes* subgenera into three main clades. (C) Phylogenetic studies incorporating various molecular datasets (notably Soghigian et al 2017 which has the most aedine taxa represented to-date) reveal increasing evidence for a three-clade hypothesis in Aedini. See Supplementary Table S1 for detailed list of taxa within each clade.

In the largest molecular phylogenetic analysis of *Aedes* species to date with more than one molecular marker, Soghigian et al. recovered three major clades of Aedini—*Psorophora* (found in the Americas, and thus not considered in Belkin 1962); a clade composed of subgenera such as *Ochlerotatus* and *Finlaya*, and the genera *Opifex* and *Haemagogus* (again, only found in the Americas); and a clade comprised of the subgenera *Aedes, Aedimorphus, Stegomyia*, and others, along with the genera *Verrallina, Armigeres*, and several others (Soghigian et al., 2017). These clades correspond well to those described by Belkin and are also quite similar to the groups of subgenera that were used as justification for elevating so many subgenera in the genus *Aedes* to generic status (Reinert et al., 2009). However, Soghigian et al. cautioned against using their molecular phylogeny for taxonomic purposes, as it drew heavily on nucleotide sequence data from GenBank, which could have been misidentified, and contained less than one sixth of all species in the tribe (although it sampled 60 subgenera and 10 genera).

When Wilkerson et al. reversed *Aedes* to its status prior to 2000, resulting in *Aedes* once again containing more than 800 species, the authors noted that the existing molecular phylogenies had sampled too few species, or the DNA fragments analyzed did not have ability to resolve the genus properly (Wilkerson et al., 2015). Since that time, numerous molecular phylogenetic studies—including studies using exome data (Soghigian et al., 2023) and whole mitochondrial genomes (Chen et al., 2024; Chu et al., 2018; da Silva e Silva et al., 2022; Ma et al., 2022) — have been released. To assess whether recent molecular phylogenies provide a clarified picture of the systematics of the genus *Aedes* when considered together, we present a review and synthesis of studies which generated current molecular phylogenies. We evaluate whether a three-clade hypothesis (Figure 1C, Table 1), based on Belkin (1962) and Soghigian et al. (2017), and supporting two clades of *Aedes* mosquitoes and a third clade for *Psorophora*, is also supported by previous phylogenetic studies. This review is intended to guide efforts to stabilize the taxonomy of *Aedes*, in light of the growing consensus emerging from the molecular phylogenetics literature.

**Table 1.**
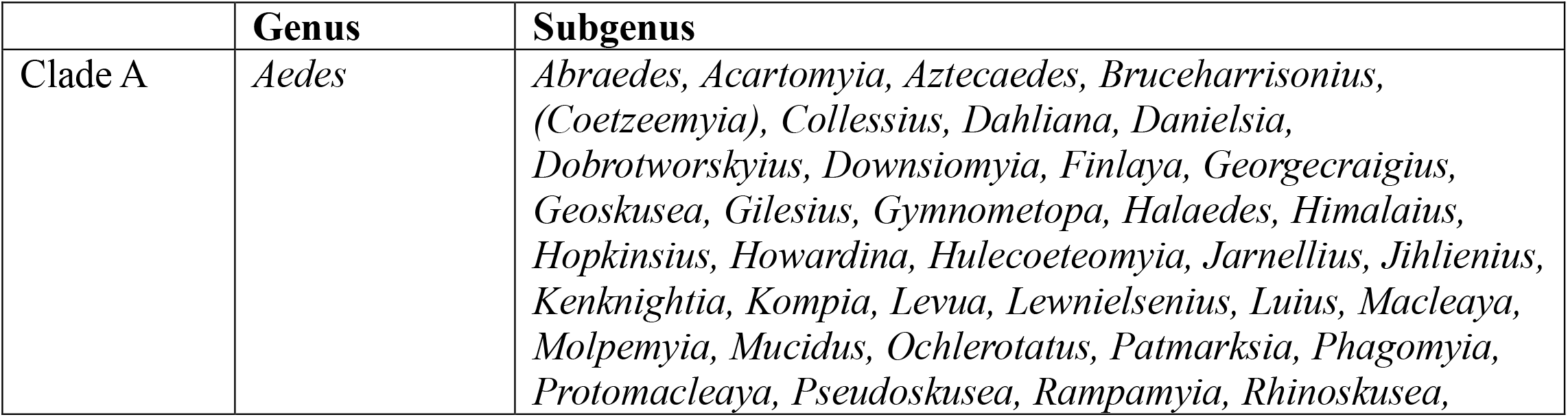
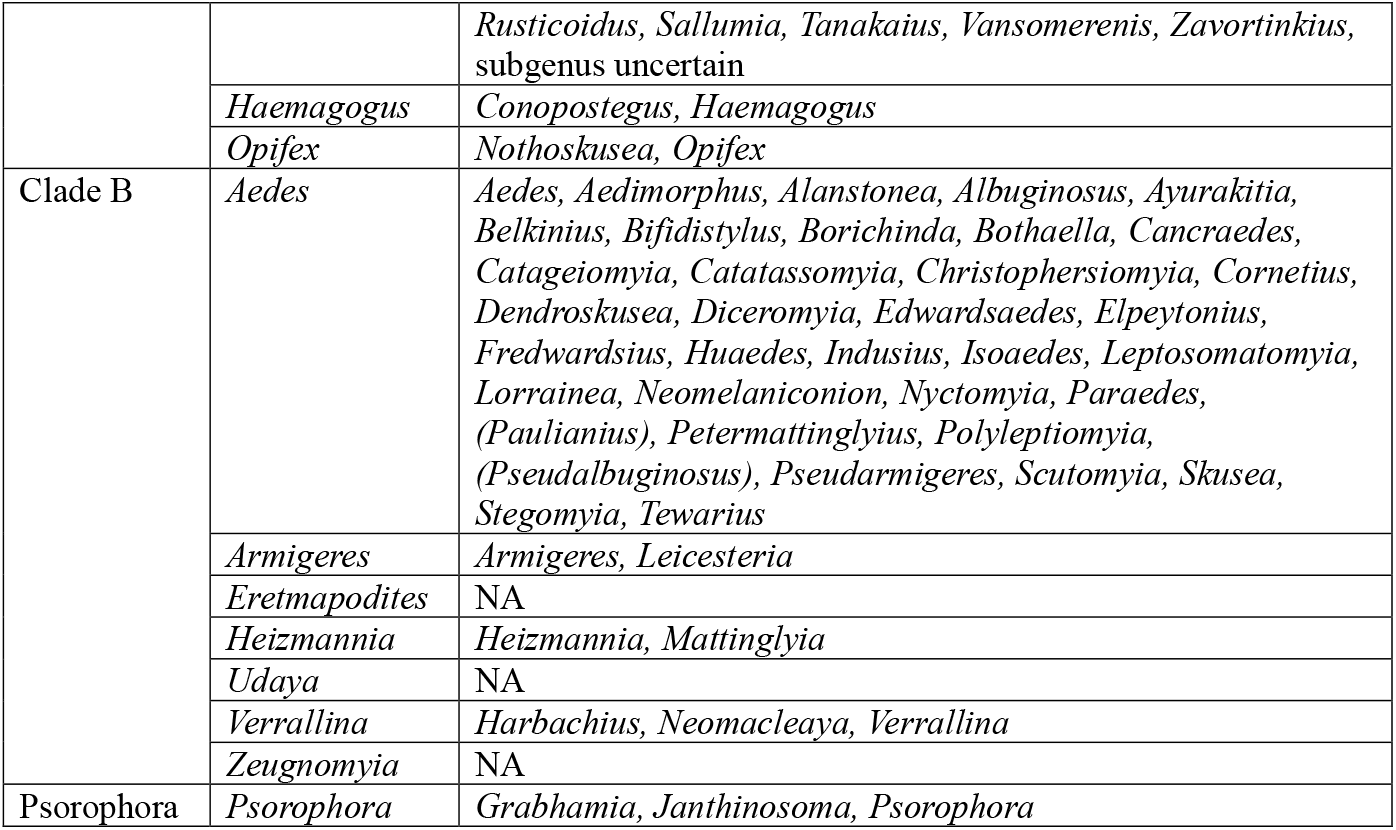
The three-clade hypothesis for tribe Aedini. Subgenus indicated with parentheses means that no morphological or molecular phylogenies have been constructed for the subgenus so far and the clade assignment is inferred. Subgenus ‘uncertain’ refers to taxa that have no current subgenus placement following Wilkerson et al. (2021) (Wilkerson et al., 2021).

## 2. Methods

We reviewed all available published studies to determine whether there was support in the literature, beyond the aforementioned studies, for three clades of the Aedini, which we term the three-clade hypothesis. The three-clade hypothesis for the tribe Aedini divides the 10 Aedini genera into three distinct clades: *Psorophora*, and two clades of *Aedes* mosquitoes, each comprised of *Aedes* subgenera and other Aedini genera (Table 1). Support for this hypothesis was first mentioned by Belkin in 1962 (Belkin, 1962) outlining Sections A and B among South Pacific mosquitoes based on morphological characters, with the notable exception that several aedine genera and subgenera, being either not yet established nor found in the South Pacific, were excluded. Following the section lettering from Belkin, we define clade A to contain Aedini genera *Opifex, Haemagogus* and *Aedes* subgenera such as *Ochlerotatus* and *Finlaya*, among many others. Clade B contains Aedini genera *Armigeres, Verrallina, Heizmannia, Zeugnomyia, Udaya, Eretmapodities*, and *Aedes* subgenera such as *Stegomyia, Aedes*, and *Aedimorphus*, among others, with *Psorophora* as an outgroup being sister to all other remaining Aedini genera.

To choose papers for our review, we conducted a literature search with combinations of keywords “Culicidae” or “Aedes”, and “phylogeny” on Google Scholar, and included studies published from 2005 onwards. We excluded studies that were unpublished (e.g. theses) and those that did not include more than one genus (or subgenus, if all mosquitoes present were *Aedes*) of the Aedini. We also excluded studies that used standard DNA barcoding markers (i.e., cytochrome oxidase subunit 1, internal transcribed spacer 2) singly, as they are inadequate for resolving deep divergences, such as those for *Aedes* where the last common ancestor is estimated at 60–100 million years ago (Soghigian et al., 2023, Soghigian et al., 2017; Zadra et al., 2021). However, we made exceptions to include phylogenies based on 18S and 28S ribosomal RNA genes as these are phylogenetically more informative (Gillespie et al., 2006). When a study presented multiple phylogenies, we selected published trees inferred from multiple genes over trees supported by one or just a few of the sampled markers. In cases where multiple genes were used in the study, but trees were analysed with single genes separately, but did not include a concatenated dataset, we considered all the trees generated in that study (Cook et al., 2005). For studies which incorporated multiple tree inference methods (maximum parsimony, maximum likelihood, Bayesian inference), we selected the phylogenetic tree indicated by the authors as the best supported result in the published paper (i.e., usually the tree with best support overall and congruent with most of the alternate trees generated).

We retrieved papers which constructed phylogenetic trees of the genus *Aedes*, using both molecular and morphological data to compare and assess whether our proposed three-clade hypothesis was reflected among published phylogenetic trees. From each study, we recorded each species and their clade membership (A, B, neither, or *Psorophora*). We then summarize the results to evaluate support for our three-clade hypothesis. We then compared the data to the phylogenetic tree in RH&K 2009 (Reinert et al., 2009) based on morphology to determine whether there is agreement between the morphological and molecular phylogenies for our three-clade hypothesis. A total of 18 publications (17 with molecular phylogenies, one with morphological phylogeny) is included in this review.

To keep the nomenclature standardized, we follow the classification of tribe Aedini from Wilkerson et al. (Wilkerson et al., 2021). As part of the process of standardizing the nomenclature, we collated a list of extant mosquito species and included the classifications of both RH&K 2009 and Wilkerson et al. 2021 (Harbach, 2025; Wilkerson et al., 2021). We have made the excel sheet accessible on a public platform (https://www.soghigian-lab.net/mosquito-species-list) to facilitate access to this information as part of a collective effort to standardize the Aedini nomenclature.

## 3 Results and Discussion

### 3.1. Challenges of resolving the phylogeny of tribe Aedini

Although numerous phylogenetic analyses have focused on mosquito species, particularly within clades of medically important species such as the Aegypti Group, relatively few studies have examined evolutionary relationships across the Aedini tribe as a whole (Yee et al., 2022). This is in part due to the size of the tribe, which includes approximately 1,300 species (Harbach, 2022), and its worldwide distribution, making comprehensive taxon sampling a logistical challenge. Among the molecular phylogenies reviewed, only one publication included representatives from all 10 aedine genera, and just three studies (17.6%) included more than 22 *Aedes* subgenera (out of a total of 79) in their phylogenetic analyses.

Aside from challenges associated with taxon sampling, there is a notable scarcity of studies that incorporate phylogenetically informative markers. We identified only 17 publications over the last 20 years that meet our criteria for including molecular phylogenies in this review. Most studies focused on only the COI barcode, a marker which is effective for species identification (Pentinsaari et al., 2016) but insufficient for resolving deeper evolutionary relationships and was thus excluded from our review. The cost and complexity of sequencing multiple genes (both mitochondrial and nuclear genes) or generating genomic data likely limited the scope of many studies, even when broad sample representation was available (Coissac et al., 2016). While elucidating the evolutionary relationships of *Aedes* was not the primary aim of most of these studies, the lack of aedine species represented in these independent studies has made it difficult for workers to come to a consensus on stabilising the classification of Aedini.

Our review approach provides a framework for evaluating genus and subgenus level placements across independent phylogenetic studies. By comparing results across studies that employ a range of phylogenetic inference methods, we assess the consistency and robustness of clade level relationships. This approach emphasizes broader evolutionary patterns within Aedini, rather than narrowly focusing on the placement of individual species, thus enabling a more integrated understanding of relationships among genera and subgenera.

### 3.2. Consensus for a three-clade hypothesis

We evaluated 22 molecular and morphological phylogenetic trees from 18 published papers over the last 20 years found overwhelming support for our proposed three-clade hypothesis (Figure 2), and with it, further evidence that *Aedes* is not monophyletic. These molecular phylogenies contained 294 aedine species from 10 genera, and we consistently found robust evidence for a three-clade hypothesis from multiple data types (i.e., single genes, multi-loci, and genomic data) and inference methods (neighbour-joining, maximum parsimony, maximum likelihood and Bayesian inference). Clade A contains *Haemagogus, Opifex*, and *Aedes* species in subgenera *Ochlerotatus* and various other subgenera. Clade B is comprised of *Armigeres, Heizmannia, Eretmapodities, Zeugnomyia, Udaya, Verrallina* and some subgenera from the genus *Aedes* – such as *Aedes, Aedimorphus* and *Stegomyia. Psorophora* forming the third clade was a sister group to the nine other Aedini genera (Figure 1, Table 1).

**Figure 2.**
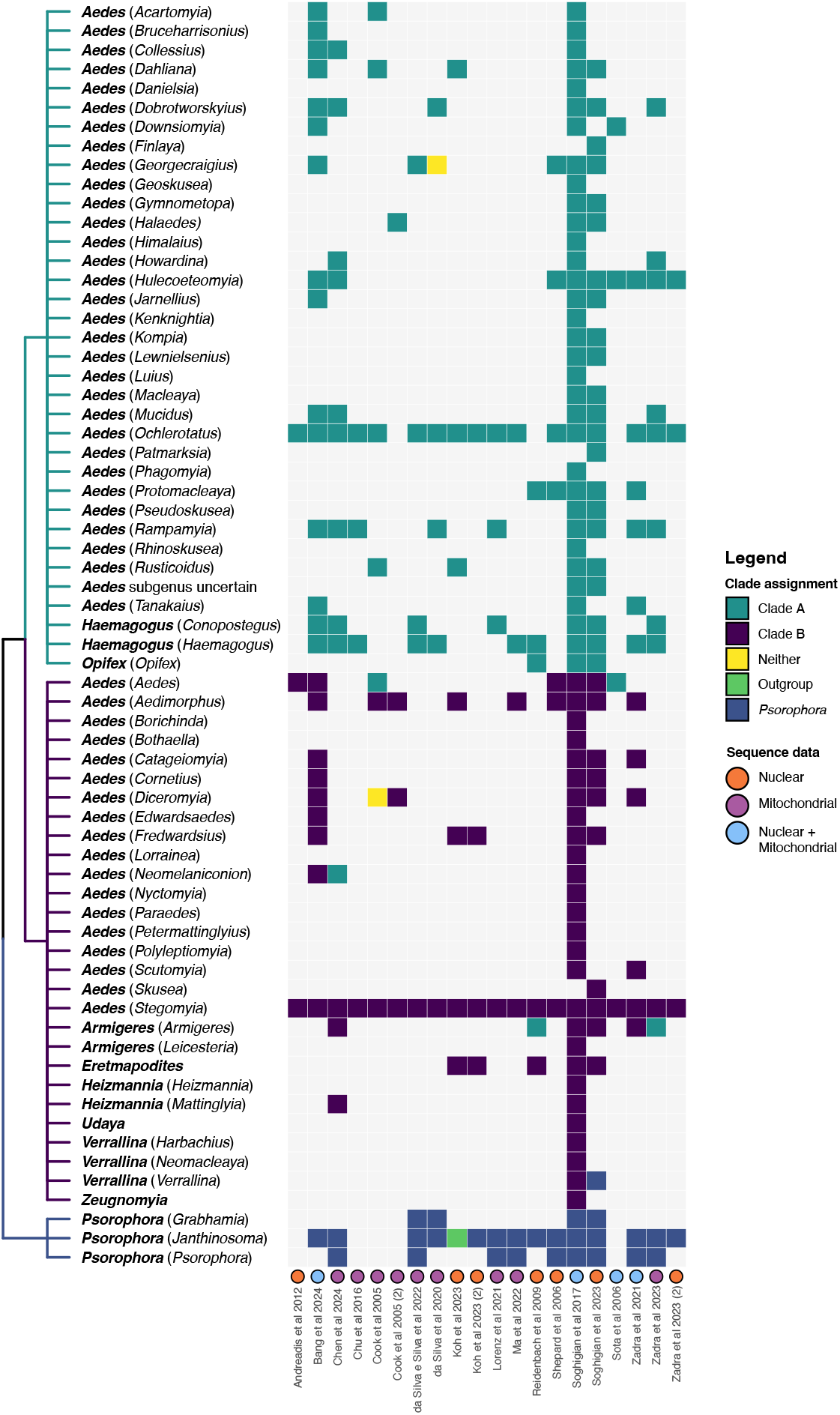
Support for the three-clade hypothesis across published molecular phylogenies. The genera and subgenera represented in all molecular phylogenies reviewed are grouped into polytomies for each clade following the three-clade hypothesis. The heatmap indicates support for the clade assignment within each sampled phylogeny.

The few conflicts with our three-clade hypothesis (Figure 2) across the phylogenies reviewed can be attributed to the lack of adequate sampling, and/or a reliance on mitochondrial genes only and poorly supported branches (Figure 2, Supplementary Table S2). In one instance, the placement of *Aedes* (*Aedes*) *esoensis* Yamada in clade A was the result of the study focusing on the subgenus *Stegomyia* and rooted the rest of the species as a separate clade (Sota and Mogi, 2006). Another species of subgenus *Aedes* (*Aedes cinereus* Theobald 1901) was placed within clade A (Cook et al., 2005); however, the node was not well-supported (bootstrap <50%) and only one barcoding marker was used in the construction of the tree. *Aedes* (*Neomelaniconion*) *lineatopennis* (Ludlow 1905) was placed in clade A in a study with eight other *Aedes* subgenera (Chen et al., 2024), but was placed within clade B in a much larger study with 47 *Aedes* subgenera and eight other *Neomelaniconion* species (Soghigian et al., 2017). *Armigeres* (*Armigeres*) *subalbatus* (Coquillet 1898) was placed in clade A in two instances (Reidenbach et al., 2009; Zadra et al., 2023), but the same species was situated in clade B in four other studies, including one where genomic data was utilised (Chen et al., 2024; Soghigian et al., 2023, 2017; Zadra et al., 2021). Two subgenera (*Diceromyia* and *Georgecraigius*) were not placed in either clade A or B because of low bootstrap support values (Cook et al., 2005) and phylogenetic inference was based only mitochondrial genes (Cook et al., 2005; da Silva et al., 2020); however, in other studies, each were resolved clades A or B. Despite the inconsistencies in clade placement of the abovementioned subgenera, when we consider the support values and studies which included other species of the same subgenera, there is overall strong evidence for the monophyly of clades A and B.

The clade placement of 49 *Aedes* subgenera and 10 aedine genera in the analysis by Reinert, Harbach, and Kitching (2009) based on morphological data alone is largely congruent with the consensus clade placement inferred from a synthesis of major findings in published molecular phylogenies (Figure 2, Supplementary figure S1). The RH&K (2009) phylogeny presented aedine species falling into five clades – 1) *Psorophora*, 2) *Aedes* (*Mucidus*), 3) the majority of species in clade A, 3) a clade consisting of *Aedes* subgenera *Sallumia, Geoskusia, Levua* and *Rhinoskusea*, and 4) clade B (Figure 1B). Where RH&K 2009 was unable to place the second or fourth clade in either clades A or B, the molecular phylogenies were able to resolve *Mucidus, Geoskusea* and *Rhinoskusea* within clade A – to date, no molecular phylogenies have sampled *Aedes* (*Sallumia*) or *Aedes* (*Levua*). Our study thus demonstrates the strength of molecular phylogenies at reconciling long-standing debates on *Aedes* phylogeny, with potential applications to taxonomy.

### 3.3. Moving forward

While our review of molecular phylogenies indicates strong support for the monophyly of three clades within the tribe Aedini, this also highlights the current paraphyletic nature of the genus *Aedes* and the need for a revision of its classification. However, we caution against a hasty reclassification of *Aedes*. Our review found that 294 aedine species are included in molecular phylogenies, only ∼22% of all aedine species described thus far. Perhaps more critically, very few studies have included representatives from many subgenera in *Aedes* – and these will be critical for constructing meaningful and monophyletic groupings of subgenera for taxonomic purposes. Molecular phylogenies have been used extensively to elucidate relationships within species groups of mosquitoes, and as we have shown here, they also consistently recover similar clades above this level, as well.

The clade placement of genera and subgenera we present here is based on available information, from molecular and morphological results – 49 *Aedes* subgenera and nine other aedine genera based on both molecular and morphological clade placement agreement, 26 *Aedes* subgenera based on morphological phylogeny alone, and one *Aedes* subgenus based on molecular phylogeny (Supplementary figure S1). At present, three subgenera (*Coetzeemyia, Paulianius*, and *Pseudalbginosus*) are unrepresented in both molecular and morphological phylogenies, two of which are monotypic. We hypothesize that *Coetzeemyia* would be in clade A as it was elevated from *Levua* (Huang et al., 2010), while *Paulianius* and *Pseudalbuginosus* are likely to be in clade B as they share morphological similarities with *Diceromyia* (Brunhes et al., 2017; Huang and Rueda, 2015) (Table 1). As detailed earlier, multiple previous attempts have been made to elevate subgenera to generic status. With the current sampling, it is largely unknown how groups of subgenera are related beyond the course three-clade level we present here. More critically, there is some evidence from molecular phylogenies (e.g. Soghigian et al. 2017) that subgenera may not be monophyletic. As such, taxonomic action along these lines at present seems premature.

Classifications and nomenclature should be “natural” (International code of zoolocial nomenclature, 1999). There is a general consensus among the scientific community that this means that naming of taxa above the species level should follow monophyletic groupings and reflect phylogenetic relationships (Vences et al., 2013). Following that, Vences et al (2013) proposed 11 Taxon Naming Criteria (TNCs) for consideration in taxonomic revisions, with monophyly, clade stability and phenotypic diagnosability being the priority TNCs (Vences et al., 2013). Also of importance, however, is that reclassification of prominent taxa with medical and economic significance should be carried out cautiously (Vences et al., 2013). In cases where non-monophyly of a taxon is known but only an incomplete phylogeny is available, a more inclusive classification to encompass the clade as a genus rather than splitting it into multiple genera should be adopted to preserve the stability of the genus for the time being (Vences et al., 2013).

Based on these criteria, it may seem that a classification based on the three clades we have found – thus splitting *Aedes* into two genera – would be sensible. However, we believe that prior to such an action, it is important that the evidence of the paraphyly in *Aedes* be demonstrated and communicated (e.g. as through this review), and that a thorough review of the morphological characters is needed to assess if the clades can be distinguished based on derived characters (i.e. synapomorphies), as such a change would also result in the demotion of long-standing genera to subgenus level (e.g. *Haemagogus, Armigeres*). Although not essential to revising the taxonomy of *Aedes*, integrating morphological characters with the results of molecular phylogenies may bring wider acceptance to taxonomic action.

We hope our results provide clarification and support that will allow development of a new standardized nomenclature for *Aedes* mosquitoes in the future. This standardization is especially critical given the role of many species as major disease vectors, where public education campaigns often use species epithets and genera, in addition to common names. As they have for so many other groups, molecular phylogenetics of *Aedes* mosquitoes have provided a strong foundation for studying evolutionary relationships particularly important in this lineage of animals, and we believe future molecular phylogenies in this group will provide a clear path towards resolving current conflicts in classification and taxonomy.

## 4.1 Acknowledgements

We would like to acknowledge the thoughtful discussions on mosquito systematics and taxonomy over the last years with T. Livdahl, R. Harbach, Y. Linton, T. Andreadis, J. Powell, C. Sither, and M. Reiskind that helped to motivate this review. Financial support for this project was provided by the University of Calgary’s PURE undergraduate research program, and we also acknowledge the support of the Natural Sciences and Engineering Research Council of Canada (NSERC), funding reference number RMS21-73779779 [Cette recherche a été financée par le Conseil de recherches en sciences naturelles et en génie du Canada (CRSNG), numéro de référence RMS21-73779779]. In addition, GM was supported by NIAID R01 AI155562 awarded to JS.

## 5.1 Author Contributions

Conceptualization: JS, BW, HY

Data Curation: CC, HQ, GM

Investigation: CC, HQ

Methodology: JS, CC, HQ

Formal Analysis: CC, HQ

Funding Acquisition: JS

Supervision: JS, BW

Writing – Original Draft Preparation: CC, HQ, JS, GM

Writing – Review and Editing: HQ, JS, GM, BW

## 6.1 Funding sources

NSERC

NIH

## References

Belkin, J.N., 1962. The mosquitoes of the South Pacific (Diptera: Culicidae). University of California Press, Berkeley and Los Angeles.

Benedict, M.Q., Levine, R.S., Hawley, W.A., Lounibos, L.P., 2007. Spread of the tiger: Global risk of invasion by the mosquito Aedes albopictus. Vector-borne and Zoonotic Diseases 7, 76–85. 10.1089/vbz.2006.0562

Brunhes, J., Boussès, P., Tantely, M.L., Kengne, P., 2017. Un nouveau genre de Culicidae (Diptera), Paulianius n. gen., avec la description de trois nouvelles espèces malgaches. Annales de la Société entomologique de France (N.S.) 53, 344–373. 10.1080/00379271.2017.1365627

Chen, D.-H., He, S.-L., Fu, W.-B., Yan, Z.-T., Hu, Y.-J., Yuan, H., Wang, M.-B., Chen, B., 2024. Mitogenome-based phylogeny of mosquitoes (Diptera: Culicidae). Insect Science 31, 599–612. 10.1111/1744-7917.13251

Chen, X.-G., Jiang, Xuanting, Gu, J., Xu, M., Wu, Y., Deng, Y., Zhang, C., Bonizzoni, M., Dermauw, W., Vontas, J., Armbruster, P., Huang, X., Yang, Y., Zhang, H., He, W., Peng, H., Liu, Y., Wu, K., Chen, J., Lirakis, M., Topalis, P., Van Leeuwen, T., Hall, A.B., Jiang, Xiaofang, Thorpe, C., Mueller, R.L., Sun, C., Waterhouse, R.M., Yan, G., Tu, Z.J., Fang, X., James, A.A., 2015. Genome sequence of the Asian Tiger mosquito, Aedes albopictus, reveals insights into its biology, genetics, and evolution. Proceedings of the National Academy of Sciences 112, E5907–E5915. 10.1073/pnas.1516410112

Chu, H., Li, Chunxiao, Guo, Xiaoxia, Zhang, Hengduan, Luo, Peng, Wu, Zhonghua, Wang, Gang, and Zhao, T., 2018. The phylogenetic relationships of known mosquito (Diptera: Culicidae) mitogenomes. Mitochondrial DNA Part A 29, 31–35. 10.1080/24701394.2016.1233533

Coissac, E., Hollingsworth, P.M., Lavergne, S., Taberlet, P., 2016. From barcodes to genomes: extending the concept of DNA barcoding. Molecular Ecology 25, 1423–1428. 10.1111/mec.13549

Cook, S., Diallo, M., Sall, A.A., Cooper, A., Holmes, E.C., 2005. Mitochondrial markers for molecular identification of Aedes mosquitoes (Diptera: Culicidae) involved in transmission of arboviral disease in West Africa. Journal of Medical Entomology 42, 19– 28. 10.1093/jmedent/42.1.19

da Silva, A.F., Machado, L.C., de Paula, M.B., da Silva Pessoa Vieira, C.J., de Morais Bronzoni, R.V., de Melo Santos, M.A.V., Wallau, G.L., 2020. Culicidae evolutionary history focusing on the Culicinae subfamily based on mitochondrial phylogenomics. Sci Rep 10, 18823. 10.1038/s41598-020-74883-3

da Silva e Silva, L.H., da Silva, F.S., Medeiros, D.B. de A., Cruz, A.C.R., da Silva, S.P., Aragão, A. de O., Dias, D.D., Sena do Nascimento, B.L., Júnior, J.W.R., Vieira, D.B.R., Monteiro, H.A. de O., Neto, J.P.N., 2022. Description of the mitogenome and phylogeny of Aedes spp. (Diptera: Culicidae) from the Amazon region. Acta Tropica 232, 106500. 10.1016/j.actatropica.2022.106500

Gillespie, J.J., Johnston, J.S., Cannone, J.J., Gutell, R.R., 2006. Characteristics of the nuclear (18S, 5.8S, 28S and 5S) and mitochondrial (12S and 16S) rRNA genes of Apis mellifera (Insecta: Hymenoptera): structure, organization, and retrotransposable elements. Insect Mol Biol 15, 657–686. 10.1111/j.1365-2583.2006.00689.x

Global Biodiversity Information Facility, 2025. Tanakaius savoryi (Bohart, 1957) [WWW Document]. URL https://www.gbif.org/species/11249369 (accessed 4.4.25).

Harbach, R.E., 2025. Valid extant species of Culicidae.

Harbach, R.E., 2022. Mosquito Taxonomic Inventory [WWW Document]. Mosquito Taxonomic Inventory. URL https://mosquito-taxonomic-inventory.myspecies.info/ (accessed 12.20.22).

Huang, Y.-M., Mathis, W.N., Wilkerson, R.C., 2010. Coetzeemyia, a new subgenus of Aedes, and a redescription of the holotype female of Aedes (Coetzeemyia) fryeri (Theobald) (Diptera: Culicidae). Zootaxa 2638. 10.11646/zootaxa.2638.1.1

Huang, Y.-M., Rueda, L.M., 2015. Pseudalbuginosus, a New Subgenus of Aedes, and a Redescription of Aedes (Pseudalbuginosus) grjebinei Hamon, Taufflieb, and Maillot (Diptera: Culicidae). went 117, 381–388. 10.4289/0013-8797.117.3.381

International code of zoolocial nomenclature, 1999. International code of zoological nomenclature. International Trust for Zoological Nomenclature, London, United Kingdom.

Inward, D., Beccaloni, G., Eggleton, P., 2007. Death of an order: a comprehensive molecular phylogenetic study confirms that termites are eusocial cockroaches. Biol Lett 3, 331–335. 10.1098/rsbl.2007.0102

Kampen, H., Werner, D., 2014. Out of the bush: the Asian bush mosquito Aedes japonicus japonicus (Theobald, 1901) (Diptera, Culicidae) becomes invasive. Parasites & Vectors 7, 59. 10.1186/1756-3305-7-59

Leta, S., Beyene, T.J., De Clercq, E.M., Amenu, K., Kraemer, M.U.G., Revie, C.W., 2018. Global risk mapping for major diseases transmitted by Aedes aegypti and Aedes albopictus. International Journal of Infectious Diseases 67, 25–35. 10.1016/j.ijid.2017.11.026

Ma, X., Wang, F., Wu, T., Li, Y., Sun, X., Wang, C., Chang, Q., 2022. First description of the mitogenome and phylogeny:Aedes vexansand Ochlerotatus caspius of the Tribe Aedini (Diptera: Culicidae). Infection, Genetics and Evolution 102, 105311. 10.1016/j.meegid.2022.105311

Misof, B., Liu, S., Meusemann, K., Peters, R.S., Donath, A., Mayer, C., Frandsen, P.B., Ware, J., Flouri, T., Beutel, R.G., Niehuis, O., Petersen, M., Izquierdo-Carrasco, F., Wappler, T., Rust, J., Aberer, A.J., Aspöck, U., Aspöck, H., Bartel, D., Blanke, A., Berger, S., Böhm, A., Buckley, T.R., Calcott, B., Chen, J., Friedrich, F., Fukui, M., Fujita, M., Greve, C., Grobe, P., Gu, S., Huang, Y., Jermiin, L.S., Kawahara, A.Y., Krogmann, L., Kubiak, M., Lanfear, R., Letsch, H., Li, Yiyuan, Li, Z., Li, J., Lu, H., Machida, R., Mashimo, Y., Kapli, P., McKenna, D.D., Meng, G., Nakagaki, Y., Navarrete-Heredia, J.L., Ott, M., Ou, Y., Pass, G., Podsiadlowski, L., Pohl, H., von Reumont, B.M., Schütte, K., Sekiya, K., Shimizu, S., Slipinski, A., Stamatakis, A., Song, W., Su, X., Szucsich, N.U., Tan, M., Tan, X., Tang, M., Tang, J., Timelthaler, G., Tomizuka, S., Trautwein, M., Tong, X., Uchifune, T., Walzl, M.G., Wiegmann, B.M., Wilbrandt, J., Wipfler, B., Wong, T.K.F., Wu, Q., Wu, G., Xie, Y., Yang, S., Yang, Q., Yeates, D.K., Yoshizawa, K., Zhang, Q., Zhang, R., Zhang, W., Zhang, Yunhui, Zhao, J., Zhou, C., Zhou, L., Ziesmann, T., Zou, S., Li, Yingrui, Xu, X., Zhang, Yong, Yang, H., Wang, Jian, Wang, Jun, Kjer, K.M., Zhou, X., 2014. Phylogenomics resolves the timing and pattern of insect evolution. Science 346, 763–767. 10.1126/science.1257570

National Center for Biotechnology Information, 2025. Tanakaius savoryi [WWW Document]. NCBI. URL https://www.ncbi.nlm.nih.gov/datasets/taxonomy/1608559/ (accessed 4.5.25).

Niehuis, O., Hartig, G., Grath, S., Pohl, H., Lehmann, J., Tafer, H., Donath, A., Krauss, V., Eisenhardt, C., Hertel, J., Petersen, M., Mayer, C., Meusemann, K., Peters, R.S., Stadler, P.F., Beutel, R.G., Bornberg-Bauer, E., McKenna, D.D., Misof, B., 2012. Genomic and morphological evidence converge to resolve the enigma of Strepsiptera. Current Biology 22, 1309–1313. 10.1016/j.cub.2012.05.018

Pentinsaari, M., Salmela, H., Mutanen, M., Roslin, T., 2016. Molecular evolution of a widely-adopted taxonomic marker (COI) across the animal tree of life. Sci Rep 6, 35275. 10.1038/srep35275

Polaszek, A., 2006. Two words colliding: resistance to changes in the scientific names of animals – Aedes vs Stegomyia. Trends in Parasitology 22, 8–9. 10.1016/j.pt.2005.11.003

Powell, J.R., Tabachnick, W.J., 2013. History of domestication and spread of Aedes aegypti - A Review. Mem. Inst. Oswaldo Cruz 108, 11–17. 10.1590/0074-0276130395

Rattanarithikul, R., Harbach, R.E., Harrison, B.A., Panthusiri, P., Coleman, R.E., Richardson, J.H., 2010. Illustrated keys to the mosquitoes of Thailand. VI. Tribe Aedini. The Southeast Asian Journal of Tropical Medicine and Public Health 41, 1–225.

Reidenbach, K.R., Cook, S., Bertone, M.A., Harbach, R.E., Wiegmann, B.M., Besansky, N.J., 2009. Phylogenetic analysis and temporal diversification of mosquitoes (Diptera: Culicidae) based on nuclear genes and morphology. BMC Evolutionary Biology 9, 298. 10.1186/1471-2148-9-298

Reinert, J., Harbach, R.E., Kitching, I.J., 2009. Phylogeny and classification of tribe Aedini (Diptera: Culicidae). Zoological Journal of the Linnean Society 157, 700–794. 10.1111/j.1096-3642.2009.00570.x

Reinert, J., Harbach, R.E., Kitching, I.J., 2004. Phylogeny and classification of Aedini (Diptera: Culicidae), based on morphological characters of all life stages. Zoological Journal of the Linnean Society 142, 289–368. 10.1111/j.1096-3642.2004.00144.x

Reisen, W.K., 2016. Update on journal policy of Aedine mosquito genera and subgenera. Journal of Medical Entomology 53, 249. 10.1093/jme/tjv169

Savage, H.M., 2005. Classification of mosquitoes in tribe Aedini (Diptera: Culicidae): Paraphylyphobia, and classification versus cladistic analysis. Journal of Medical Entomology 42, 923–927. 10.1093/jmedent/42.6.923

Soghigian, J., Andreadis, T.G., Livdahl, T.P., 2017. From ground pools to treeholes: convergent evolution of habitat and phenotype in Aedes mosquitoes. BMC Evolutionary Biology 17, 262. 10.1186/s12862-017-1092-y

Soghigian, J., Sither, C., Justi, S.A., Morinaga, G., Cassel, B.K., Vitek, C.J., Livdahl, T., Xia, S., Gloria-Soria, A., Powell, J.R., Zavortink, T., Hardy, C.M., Burkett-Cadena, N.D., Reeves, L.E., Wilkerson, R.C., Dunn, R.R., Yeates, D.K., Sallum, M.A., Byrd, B.D., Trautwein, M.D., Linton, Y.-M., Reiskind, M.H., Wiegmann, B.M., 2023. Phylogenomics reveals the history of host use in mosquitoes. Nature Communications 14, 6252. 10.1038/s41467-023-41764-y

Song, N., Wang, M.-M., Huang, W.-C., Wu, Z.-Y., Shao, R., Yin, X.-M., 2024. Phylogeny and evolution of hemipteran insects based on expanded genomic and transcriptomic data. BMC Biol 22, 190. 10.1186/s12915-024-01991-1

Sota, T., Mogi, M., 2006. Origin of pitcher plant mosquitoes in Aedes (Stegomyia): A molecular phylogenetic analysis using mitochondrial and nuclear gene sequences. Journal of Medical Entomology 43, 795–800. 10.1093/jmedent/43.5.795

Thomas, J.A., Trueman, J.W.H., Rambaut, A., Welch, J.J., 2013. Relaxed phylogenetics and the Palaeoptera problem: Resolving deep ancestral splits in the insect phylogeny. Systematic Biology 62, 285–297. 10.1093/sysbio/sys093

Vasilikopoulos, A., Misof, B., Meusemann, K., Lieberz, D., Flouri, T., Beutel, R.G., Niehuis, O., Wappler, T., Rust, J., Peters, R.S., Donath, A., Podsiadlowski, L., Mayer, C., Bartel, D., Böhm, A., Liu, S., Kapli, P., Greve, C., Jepson, J.E., Liu, X., Zhou, X., Aspöck, H., Aspöck, U., 2020. An integrative phylogenomic approach to elucidate the evolutionary history and divergence times of Neuropterida (Insecta: Holometabola). BMC Evol Biol 20, 64. 10.1186/s12862-020-01631-6

Vences, M., Guayasamin, J.M., Miralles, A., Riva, I.D.L., 2013. To name or not to name: Criteria to promote economy of change in Linnaean classification schemes. Zootaxa 3636, 201– 244. 10.11646/zootaxa.3636.2.1

Waterhouse, R.M., Wyder, S., Zdobnov, E.M., 2008. The Aedes aegypti genome: a comparative perspective. Insect Molecular Biology 17, 1–8. 10.1111/j.1365-2583.2008.00772.x

Wilkerson, R.C., Linton, Y.-M., Fonseca, D.M., Schultz, T.R., Price, D.C., Strickman, D.A., 2015. Making mosquito taxonomy useful: A stable classification of tribe Aedini that balances utility with current knowledge of evolutionary relationships. PLOS ONE 10, e0133602. 10.1371/journal.pone.0133602

Wilkerson, R.C., Linton, Y.-M., Strickman, D.A., 2021. Mosquitoes of the World. Johns Hopkins University Press.

Yee, D.A., Dean Bermond, C., Reyes-Torres, L.J., Fijman, N.S., Scavo, N.A., Nelsen, J., Yee, S.H., 2022. Robust network stability of mosquitoes and human pathogens of medical importance. Parasites & Vectors 15, 216. 10.1186/s13071-022-05333-4

Zadra, N., Rizzoli, A., Rota-Stabelli, O., 2021. Chronological incongruences between mitochondrial and nuclear phylogenies of Aedes mosquitoes. Life 11, 181. 10.3390/life11030181

Zadra, N., Tatti, A., Silverj, A., Piccinno, R., Devilliers, J., Lewis, C., Arnoldi, D., Montarsi, F., Escuer, P., Fusco, G., De Sanctis, V., Feuda, R., Sánchez-Gracia, A., Rizzoli, A., Rota-Stabelli, O., 2023. Shallow whole-genome sequencing of Aedes japonicus and Aedes koreicus from Italy and an updated picture of their evolution based on mitogenomics and barcoding. Insects 14, 904. 10.3390/insects14120904

