## Supplementary figures and images for "A Way Through the Trees: Molecular Phylogenies Consistently Recover Two Clades of *Aedes* Mosquitoes"

### Supplemental Figure 1

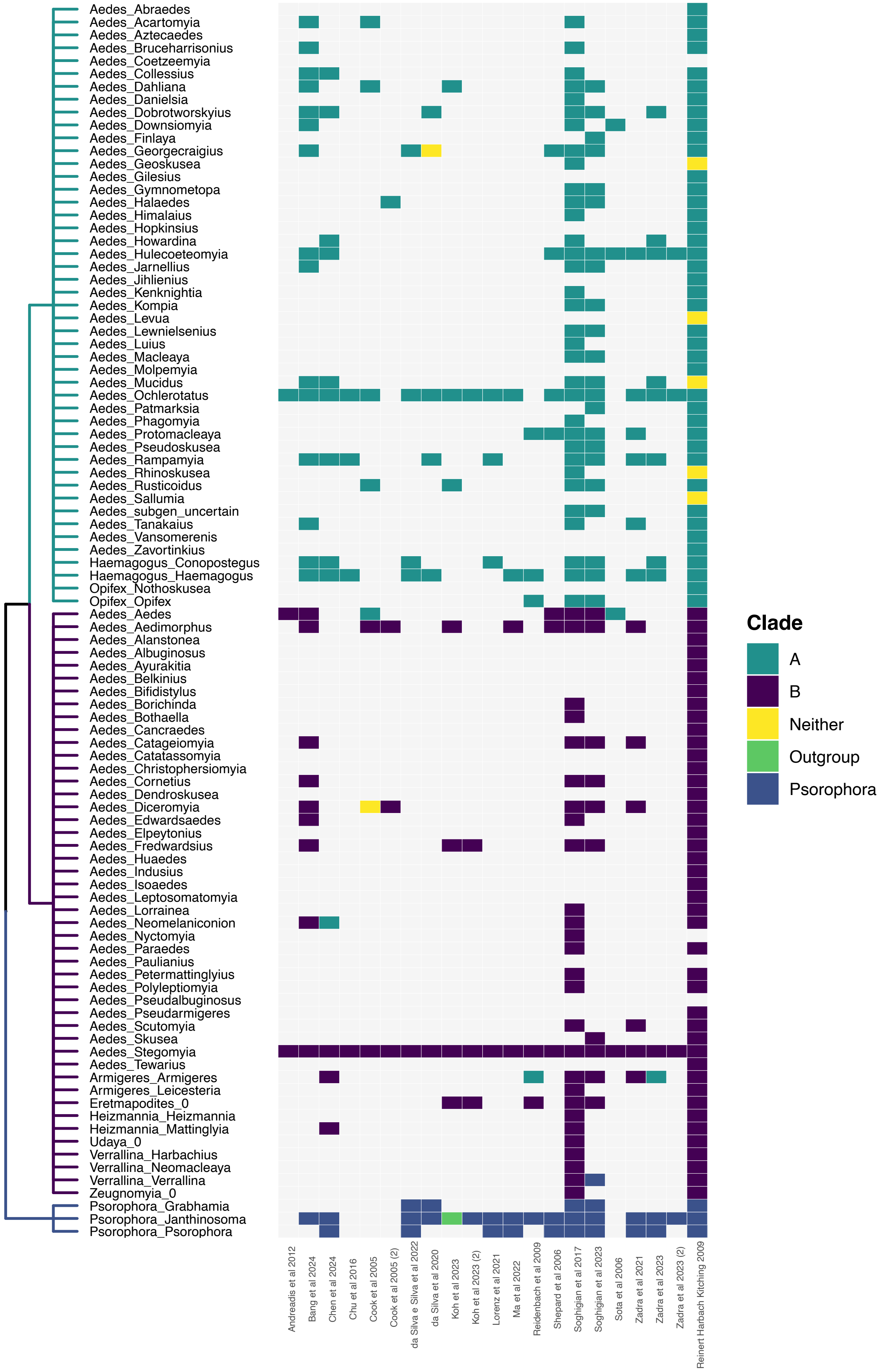
